# Model-based analysis of polymorphisms in an enhancer reveals cis-regulatory mechanisms

**DOI:** 10.1101/2020.02.07.939264

**Authors:** F Khajouei, N Samper, NJ Djabrayan, B Lunt, G Jiménez, S Sinha

**Affiliations:** University of Illinois Urbana-Champaign; Institució Catalana de Recerca i Estudis Avançats; Princeton University; University of Illinois at Urbana Champaign

**Keywords:** gene expression, enhancer, sequence analysis, ensemble of models, polymorphisms, regulatory variants

## Abstract

It is challenging to predict the impact of small genetic changes such as single nucleotide polymorphisms on gene expression, since mechanisms involved in gene regulation and their cis-regulatory encoding are not well-understood. Recent studies have attempted to predict the functional impact of non-coding variants based on available knowledge of cis-regulatory encoding, e.g., transcription factor (TF) motifs. In this work, we explore the relationship between regulatory variants and cis-regulatory encoding from the opposite angle, using the former to inform the latter. We employ sequence-to-expression modeling to resolve ambiguities regarding gene regulatory mechanisms using information about effects of single nucleotide variations in an enhancer. We demonstrate our methodology using a well-studied enhancer of the developmental gene *intermediate neuroblasts defective* (ind) in *D. melanogaster*. We first trained the thermodynamics-based model GEMSTAT to relate the neuroectodermal expression pattern of ind to its enhancer’s sequence, and constructed an ensemble of models that represent different parameter settings consistent with available data for this gene. We then predicted the effects of every possible single nucleotide variation within this enhancer, and compared these to SNP data recorded in the Drosophila Genome Reference Panel. We chose specific SNPs for which different models in the ensemble made conflicting predictions, and tested their effect in vivo. These experiments narrowed in on one mechanistic model as capable of explaining the observed effects. We further confirmed the generalizability of this model to orthologous enhancers and other related developmental enhancers. In conclusion, mechanistic models of cis-regulatory function not only help make specific predictions of variant impact, they may also be learned more accurately using data on variants.

**STATEMENT OF SIGNIFICANCE:** A central issue in analyzing variations in the non-coding genome is to interpret their functional impact, and their connections to phenotype differences and disease etiology. Machine learning methods based on statistical modeling have been developed to associate genetic variants to expression changes. However, associations predicted by these models may not be functionally relevant, despite being statisticaly significant. We describe how mathematical modeling of gene expression can be employed to systematically study the non-coding sequence and its relationship to gene expression. We demonstrate our method in a well studied developmental enhancer of the fruitfly. We establish the efficacy of mathematical models in combination with the polymorphism data to reveal new mechanistic insights.

## INTRODUCTION

Rapid advances in sequencing technologies promise to lead us to a new era of personalized diagnosis and treatment based on genomics (1). One of the most pressing challenges in this era will be to uncover the cellular and physiological impacts of DNA variations (2, 3), especially non-coding variations. A vast majority of variations associated with complex traits and common diseases fall in non-coding regions of the genome (4–6), and likely impact gene regulatory functions (7, 8). While statistical genetics approaches have proven invaluable in short-listing such variants in specific disease contexts (9), to elucidate the mechanistic basis of these variants one needs a detailed quantitative understanding of regulatory sequence function that is often beyond state-of-the-art (10).

The non-coding genome exhibits a hierarchical organization of structural and functional units, including large topologically associating domains or TADs at the megabasepair-scale (11), accessible regions and enhancers at the kilobasepair-scale (12, 13) and transcription factor (TF) binding sites at the basepair-scale. It is believed that a common mechanism of variant impact on cellular function is by affecting TF binding site strength, and consequently the gene expression level driven by an enhancer (14, 15). Thus, to investigate such mechanisms we need a precise quantitative method to predict expression level from enhancer sequence, i.e., a “sequence-to-expression” model (16–22); such methods must be sensitive enough to predict the regulatory effect of relatively minor changes in enhancer sequence, as is often the case with individual variations.

Sequence-to-expression models have been proposed in the literature to address the above need (23). These are mathematical models, based on biophysical principles (24) or machine learning concepts (25, 26), that map an enhancer’s sequence, optionally along with additional contextual information such as cellular concentrations and DNA-binding preferences of TFs, to the expression level driven by that enhancer. These models formalize what is known about a gene’s regulatory mechanisms encoded in enhancers, and have proven capable, in some cases, of predicting the effects of minor sequence differences such as mutagenesis of entire binding sites (19, 27, 28). However, when using these models one is faced with a trade-off in predictive accuracy: one fits the model to many different enhancers (22) if one wishes to capture broad regulatory mechanisms, but the resulting models are not capable of predicting the effect of minor changes such as single nucleotide variations. To achieve this latter capability, one typically fits the model to fewer, more closely related enhancers (27), but this results in under-constrained models and parameter uncertainty (29). The result is not a unique trained model but an ensemble of models, representing distinct mechanistic explanations of the data, and thus an ensemble of predictions about the effect of the same sequence mutation. If additional information becomes available about the true effect of an enhancer mutation on gene expression, that information may be found to be consistent with only a subset of the current ensemble of models and thus allow us to filter the ensemble and reduce our uncertainty about parameters, consequently increasing our confidence about cis-regulatory encoding of the enhancer. This is the key insight we pursue in this work.

We first used a thermodynamics-based modeling framework to fit an ensemble of models that relate the expression pattern of the gene *intermediate neurons defective* (ind) to the known enhancer of the gene. Thermodynamics based models are among the most successful genre of quantitative models for the sequence-to-expression relationship (19, 24), and formalize the enhancer’s cis-regulatory encoding through model parameters that are fit to the available data. We had previously shown how ensemble modeling of the thermodynamics-based GEMSTAT model (27, 29) can provide useful insights about the *ind* enhancer. Here, we used the ensemble of models to predict the effect of each single nucleotide mutation in the enhancer, and used our previously published probabilistic framework to identify mutations with high expected impact and/or high variance in predicted impact. We experimentally tested the effect of such mutations, using transgenic reporter assays in fruitflies, and used the resulting additional information to reduce the ensemble of models to a single tightly clustered set of models that represent a unique mechanistic explanation of the enhancer’s function. We then showed that the resulting model is indeed supported by additional data not used in the modeling, e.g., it provides better fits to unseen enhancers related to the *ind* enhancer. Our work attempts for forge a path forward towards deeper mechanistic understanding of the cis-regulatory ‘code’ (30) and its use in predicting the impact of single nucleotide variants (31).

## METHODS

### Model training

We set up the GEMSTAT model with 13 different parameters as in (27, 29) and trained the model to relate the wild-type expression profile of the *ind* enhancer to the concentration profiles of the five TFs DL, ZLD, SNA, VND, CIC. GEMSTAT uses the following parameters: *K*_*TF*_ parameter indicating TF-DNA binding (one for each TF), *α*_*TF*_ parameter to capture the TF’s effect on transcription rate (one for each TF), and the parameter *q*_*BTM*_ for the basal transcriptional rate. Based on previous experimental studies (32), DL and ZLD work cooperatively and this was modeled through the cooperativity parameter *w*_*DL*–*Zld*_. Previously reported reduction in CIC-DNA binding by locally activated ERK is modeled using the *Cic*_*att*_ parameter, as explained in (27). Henceforth, we refer to any setting of values for the above 13 parameters as a model. We uniformly sampled the defined range of each parameter and measured the SSE score between the wild-type and predicted expression profiles. We used a loose threshold to filter for models with good fits (SSE < 0.15), used these models as starting points for optimization (following GEMSTAT’s in-built optimization) and then selected optimized models that meet a strict threshold (SSE < 0.05). The result is an ensemble of 5237 models with distinct parameter settings that produce high quality fits to the expression profile of the gene. This is called the “wild-type ensemble”, as it was trained solely on the wild-type expression profile.

Additionally, we utilized data from six different perturbation experiments pertaining to *ind* gene expression in the same developmental stage as above. These biological perturbation experiments included (**i)** ‘DL 1 site’ (33), where mutagenesis of the strongest DL site results in no significant change of *ind* expression, (**ii)** ‘DL 3 sites’ (34), where mutagenesis of three DL sites results in greatly diminished *ind* expression, (**iii)** ‘ZLD sites’ (27), where *ind* peak expression reduces to ∼50% of wild-type levels upon mutagenesis of four ZLD sites, (**iv)** ‘SNA KO’ (18), where knock-out of SNA results in no significant change, (**v)** ‘VND KO’ (35), where knock-out of VND leads to ventral expansion of *ind* expression, and (**vi)** ‘CIC site mut.’ (36), where CIC site mutagenesis leads to dorsal de-repression. Each of the 5237 models in the wild-type ensemble was used to predict the effect of each of these six perturbation experiments, and only those models whose predictions were consistent with data for at least 5 out of the 6 perturbations were retained (Supplementary figure 7). Only 12 of the 5237 examined models met this requirement and none of these makes predictions consistent with all six perturbation experiments; in particular, no model was able to explain the ‘DL 3 site’ and ‘ZLD sites’ perturbations simultaneously (Supplementary figure 7G). To obtain a larger ensemble of models similar to these twelve, we sampled 10000 points in the parameter space around each of the 12 models and optimized the parameters to fit the wild-type expression profile of *ind*, using the sampled points for initialization. We retained from among the resulting models only those whose predictions were consistent with the perturbation experiments by the above-mentioned criterion. The retained models cluster into 18 different groups (as per method noted in the next paragraph), and we sampled 100 models from each group to obtain an ensemble of 1800 models in total. This collection of models is referred to as the “filtered ensemble”.

### Systematic prediction of SNP effects using ensemble of models

We then followed the procedure in our previous work (29) to first cluster all models in an ensemble and then construct a probability distribution over the models such that each cluster (or group) of models has the same overall probability. We then computed the average predicted effect of every possible single nucleotide mutations in the *ind* enhancer. The effect of a mutation was computed by comparing a model’s predicted expression profile for the wild-type enhancer to that for a mutated enhancer (carrying the specific mutation), and recording the sum-of-squared-errors (SSE) between the two profiles. We repeated this procedure for every model in the ensemble and computed the mean (as well as variance) of predicted effects, over the above-mentioned probability distribution over the ensemble. Such ensemble averages were separately computed for the wild-type ensemble, the filtered ensemble as well as for each cluster of models within the latter.

### Statistical testing of compensatory effects of SNPs

We generated 2000 synthetic genotypes representing the *ind* enhancer that exhibit polymorphisms at the 70 SNP positions in the DGRP population, at the same allele frequencies as in the population. For this, we first represented the DGRP genotypes as a genotype matrix where the rows represent SNPs, columns represent lines and each cell in the matrix is 0 or 1 depending if a line carries the SNP. We permuted this matrix while preserving the sum of each row as well as each column. The permutation process selects at random two rows and two columns such that each row and each column of the resulting 2 x 2 matrix has exactly one ‘0’ and one ‘1’, and swaps its rows, and the repeats this operation 1000 times. At the end, a column is chosen at random from the resulting permuted matrix and represents a sampled genotype. For each sampled genotype, we made model-based predictions of the effect (SSE) of all mutations present in that genotype as well as the effects of each mutation individually, and deemed the genotype as exhibiting compensatory mutation if the effect of all mutations together was less than the strongest among individual mutation effects. We compared the number of genotypes with compensatory mutation (964 of 2000) to the corresponding number in the DGRP (168 of 205), using a Fisher’s exact test.

### In vivo reporter assays

*ind*^*1.4*^ enhancer constructs were prepared using as a template the wild-type version of the enhancer present in DGRP line *RAL-821*. Mutagenized enhancers carrying changes at positions 1198, 309 and 309 + 324 (“construct1”, “construct2”, “construct3” repesctively, see Results) were generated by recombinant PCR and sequenced to confirm their integrity. The final transgenes were assembled in the *placZ-attB* vector (37) and integrated at chromosomal position 86F via ΦC31-mediated germ-line transformation (38).

### Reagents

Primary: Sheep anti DIG Roche Cat# 11333089001; RRID: AB_514496

DAPI: ThermoFisher Scientific Cat# D1306; RRID: AB_2629482

Secondary: Alexafluor Donkey Anti Sheep 555 Catalog # A-21436 RRID:AB_2535857

### Animal Care

Adult flies were matured at 25°C on fresh yeast paste for at least three and at most 14 days to collect embryos. All embryos used in this study were at developmental stages prior to gastrulation, which precluded the determination of the sex of embryos. All embryos were grown on apple juice plates at 25°C. All fly stocks were maintained by standard methods at 25°C, and were grown on a standard cornmeal, molasses, and yeast media. Fly media recipe: water (1726 mL), agar (11 g), potassium sodium tartrate (12 g), calcium chloride dihydrate (1 g), sucrose (43.35 g), dextrose (86.65 g), yeast (44 g), cornmeal (105 g), propionic acid (10 mL). Prepare as follows: measure water into kettle, mix in agar and bring to a boil to melt agar. Slowly add potassium sodium tartrate, calcium chloride dihydrate, sucrose, and dextrose, stirring as you add. Bring to a boil. Mix yeast and cornmeal with a little water, and add to kettle and stir. Boil 2 mins. Cool to 80°C and add propionic acid. Stir very well.

### FISH

Embryos from lines expressing lacZ under the control either WT or variant promoters were collected and allowed to develop to late embryonic stage 6. Embryos were then dechorionated in 50% bleach and fixed in 4% formaldehyde in PBS for 20 min. Embryos were then incubated in 90% xylene for 1 h and then treated with 80% acetone for 10 min at −20°C. Next, embryos were hybridized overnight at 60°C with antisense probes against lacZ mRNA labeled with digoxigenin (DIG) (1:25). Embryos with labeled probes were visualized using standard immunofluorescence technique. The following primary antibodies were used in this study: sheep anti-DIG (Roche; 1:200).

### Imaging

Embryos were imaged using a Nikon A1R-Si confocal microscope. Images were processed in FIJI to adjust levels and crop.

### Model predictions for *rho* enhancer

These models had eight parameters, four of which (*K*_*DL*_, *α*_*DL*_, *K*_*SNA*_, *α*_*SNA*_) were shared with models for the *ind* enhancer and were kept fixed at values trained on the *ind* data, while the other four (*K*_*TWI*_, *α*_*TWI*_, the cooperativity parameter *w*_*DL*–*TWI*_, and *q*_*BTM*_) were trained on *rho* data set of Sayal et al. (19).

## RESULTS

### Expression data support diverse mechanistic models of *ind* enhancer function

Our first goal was to train a model capable of predicting the impact of enhancer sequence changes on gene expression. For this, we considered various models in the literature that can predict gene expression profile from an enhancer’s sequence and information about transcription factor (TF) concentrations and their DNA binding preferences (motifs) (17–19, 39). We chose to work with one such model, called GEMSTAT (22), which was previously reported by us and successfully used to model several developmental enhancers of Drosophila (27, 40). GEMSTAT uses a statistical thermodynamics formulation to capture the molecular interactions between TFs and DNA and their quantitative impact on transcription rate. It uses two tunable biophysical parameters for each TF: one parameter that represents activation/repression strength and the other related to DNA-binding strength of the TF at its optimal site. It has one additional global parameter corresponding to the basal transcriptional machinery and optional parameters for cooperativity between specific pairs of TFs. GEMSTAT uses available data – enhancer sequence(s), expression levels, TF concentrations and TF binding specificities or ‘motifs’ – to find optimal values for its free parameters (usually ∼10-20 parameters, representing 5-10 TFs), and in some cases this procedure is known to result in locally but not globally optimal parameter values. We addressed this problem in recent work (29) by generating and reasoning with an *ensemble* of model parameters that fit the data, rather than determining a single parameter setting that maximizes the ‘goodness-of-fit’.

In the current work, we first modeled the expression data available for the ‘intermediate neuroblasts defective’ (*ind*) gene in Drosophila melanogaster, following our previous work (29). The enhancer for *ind* is well characterized, and its regulators are well-studied. There are two activators: Dorsal (Dl) and Zelda (Zld) and three repressors: Snail (Sna), Ventral neuroblasts defective (Vnd) and Capicua (Cic). The expression data includes the spatial pattern of *ind* gene and its TFs in the blastoderm stage of embryonic development, as a 1-dimensional profile along the dorso-ventral (D/V) axis (Figure 1A). By optimizing parameters of the GEMSTAT model through a comprehensive grid-search, we obtained an ensemble (‘wild-type ensemble’, see Methods) of 5237 models that produce close fits to the wild-type expression pattern (Figure 1B). (Each model is a distinct setting of tunable parameters, see Figure 1C.) We next challenged the ensemble of models with published data (27, 32–35) on *ind* gene expression under various perturbation conditions such as mutagenesis of one or more sites of a TF or knockout of a TF. Discarding models in the ensemble whose predictions were inconsistent with results from two or more of these six perturbation experiments, and performing deeper sampling and optimization around the remaining models, we constructed a new ensemble of models, henceforth called the ‘filtered ensemble’ that reasonably capture *ind* expression in wild-type as well as perturbation experiments (see Methods). Closer examination of the filtered ensemble and clustering of the parameter vectors revealed multiple groups of models in the ensemble (Figure 1D). The existence of several distinct groups of models is also clear upon visual inspection of a lower-dimensional projection of the parameter vectors using principal components analysis (Figure 1E). The predictions of each group of models in each of the perturbation conditions are shown in Figures 1F-K, along with the true expression profiles in those conditions. We noted that all groups of models were consistent with the four perturbation experiments represented by Figures 1F-I, and were additionally consistent with the experiments represented by either Figure 1J or Figure 1K.

**Figure 1.**
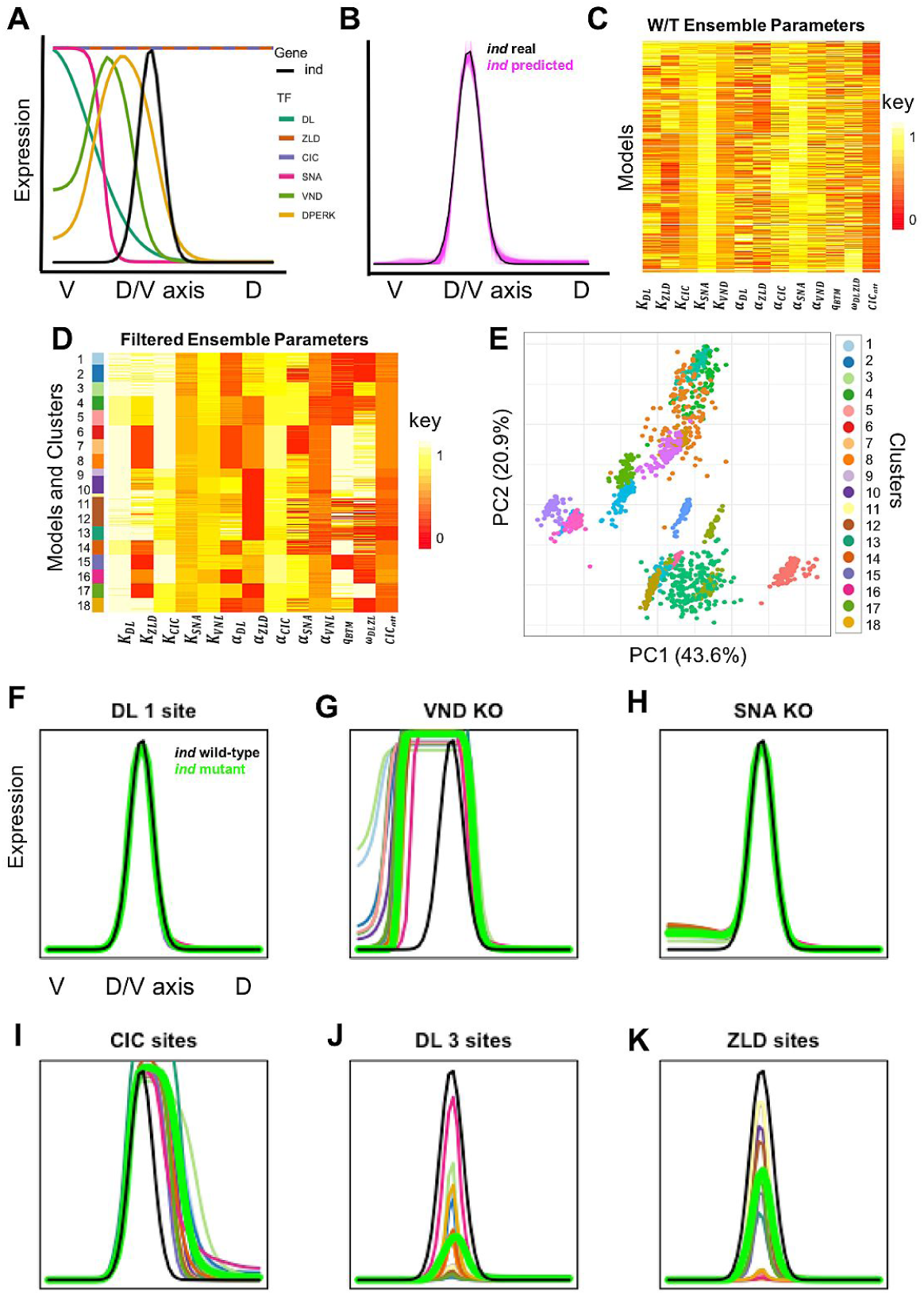
The wild-type ensemble of models combined with the perturbation experiments produce accurate predictions on ind expressions with distinct groups of models with respect to distribution of parameters in the ensemble. **(A)** The expression domain for TFs and the ‘*ind’* gene is shown along the Dorsal-ventral domain. The x-axis represents ventral (left) to dorsal (right) end of the D/V axis and the y-axis is the expression value from no expression to the maximum observed expression for each gene or TF, on a scale of 0 to 1. **(B)** Predicted *ind* expression (magenta) from all models optimized to fit wild-type data (black) and the perturbations. Each pink line shows the prediction of a single model in the ensemble **(C)** Heatmap of model parameters in B, that fit the wild-type *ind*. Each row is a model in the ensemble and each column corresponds to a parameter for the model. Each parameter is scaled to the range of 0 to 1. The K parameter for all TFs and *α* parameter of repressors are in logarithmic scale and the *α* parameter of activators, cooperativity and *q*_*BTM*_ are in linear scale. **(D**) Similar to C, heatmap of the filtered ensemble of models with sidebar colors are different groups of models that cluster together. **(E)** We projected the 13-dimensional parameter space of the filtered ensemble into the first two principal components. The scatter plot shows different groups of models, cluster with respect to their groups. Models with similar parameters are clustered together and we can observe they are separate from each other. Colors are similar to D sidebar. **(F)** No change is observed in the expression when the strongest DL site is mutated. The green shows the experimental data (Expected expression profile in D) the predicted expression from different groups of models in D. the colors are similar to the sidebar colors in D and E for different groups of models **(G)** *ind* expression expands ventrally in VND knockout. **(H)** The expression of *ind* is not changed in SNA knockout experiment. **(I)** *ind* expression expands dorsally when two sites of CIC is mutated. **(J)** Peak *ind* expression is reduced by 65% after 3 DL sites are mutated. **(K)** Peak *ind* expression is reduced by half upon mutations in ZLD binding sites.

Figure 1E reveals uncertainty about mechanisms underlying the gene’s regulation, even after subjecting the model to data from the several experimental conditions noted above. Each group or cluster of models represents a distinct hypothesis about the regulatory mechanisms underlying *ind* regulation and further information is necessary to narrow down the possible mechanisms. We looked to polymorphism data from a population of *D. melanogaster* lines (41) for such information, as described next.

### Model-based analysis of polymorphisms in the *ind* enhancer

We assessed all possible single nucleotide mutations in the *ind* enhancer (length 1416 bp, Figure 2A) using the filtered ensemble as follows: for every position in the enhancer, for every possible mutation at that position, we predicted the effect of the specific mutation using each model in the ensemble. We measured the magnitude of the predicted effect as the ‘sum of the squared errors’ (SSE) between model-predicted expression profile of the wild-type enhancer and predicted profile of the wild-type enhancer modified by that particular mutation. We then summarized the effect (SSE) of the mutation as predicted by all models in the ensemble, using a probability distribution over the filtered ensemble constructed as in (29). The predicted effect of a mutation is defined as the SSE averaged over the ensemble. Figure 2B shows these predicted effects for every position of the *ind* enhancer, aligned with the position and strength of each TF binding site in the wild-type sequence (Figure 2A). (For each position, only the mutation with greatest predicted effect is shown; see Supplementary Figure 1 A-B for examples of such effects.) Following the same procedure, we also computed variance of the predicted effect among models in the ensemble (Figure 2C). The ‘heatmaps’ in Figure 2D depict the effect of all possible mutations within three specific transcription factor binding sites (TFBS) that are strong matches to their respective motifs and harbor mutations with relatively large predicted effects. Most models in the filtered ensemble predict that the gene expression should change significantly upon mutating at least one position within these strong binding sites (Figure 2D). Predictions were not entirely consistent among models though, and different models vary in predicted extent of change. For instance, Supplementary Figures 1 A, C reveal discrepancies among the predicted effects of a mutation in a VND site, as predicted by different models in the ensemble.

**Figure 2.**
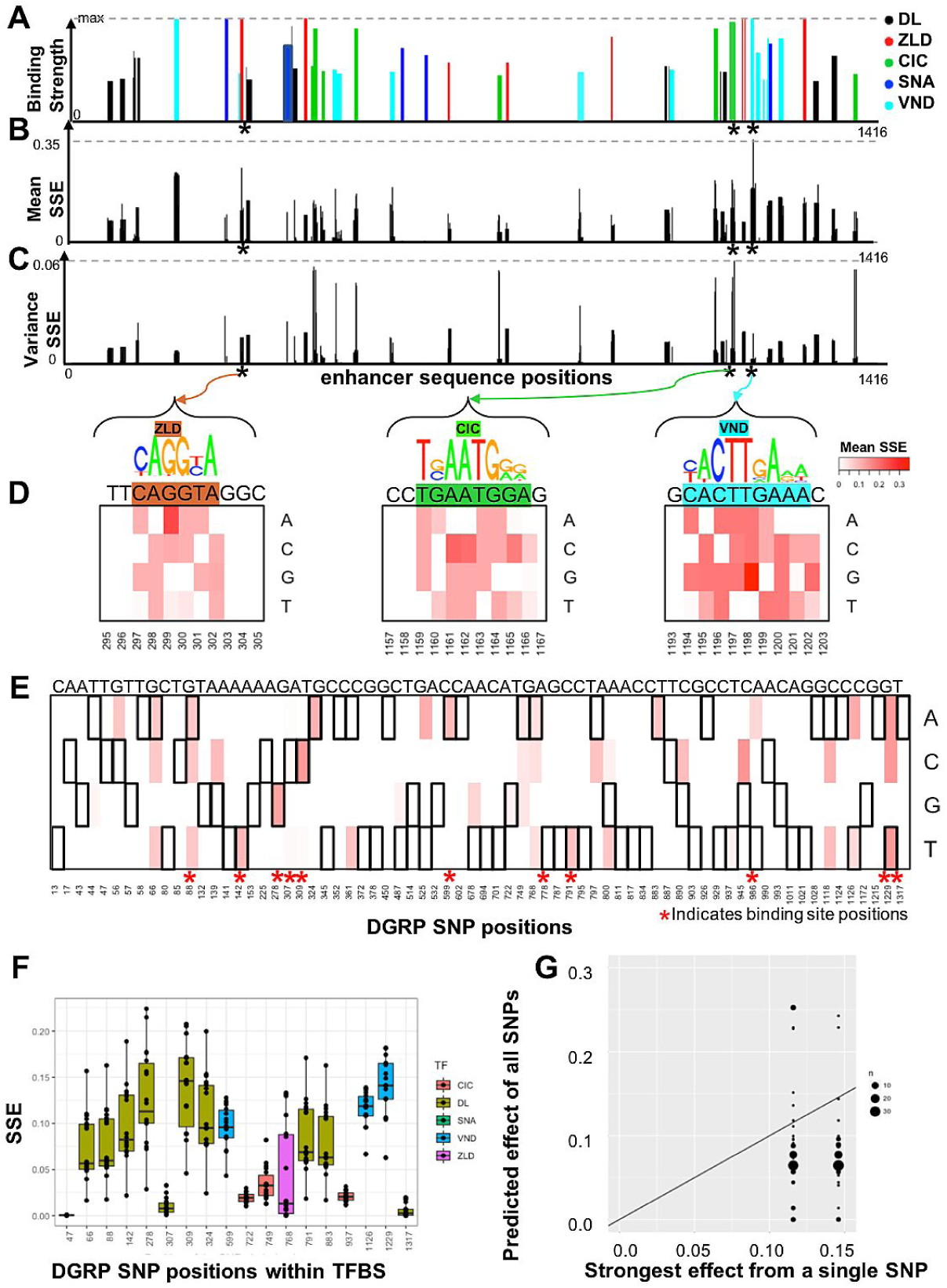
Model-based analysis of polymorphism in the ind enhancer points out to specific positions in the enhancer. **(A)** Binding sites on the ind enhancer. The x-axis shows positions in the enhancer and the y-axis is the binding strength of each TF. Binding strength is calculated from the binding energy of the motifs for each TF. **(B)** At each position of the enhancer, we introduce a single mutation (3 possible changes at each position) then we measure the SSE score between the wild-type expression profile and the mutant expression predicted by a model. We averaged the predicted SSE score across different models in the ensemble and reported the maximum score between the three average expected changes **(C)** Similar to part B, we reported maximum of the standard deviations of the expected SSE score (for the 3 possible changes of the base pair) at each position **(D)** The heatmaps show the expected SSE score for each mutation at each position. We selected the top average SSE in all enhancer positions for three TFs and showed the associated TF binding motif at the locations. Stars indicate the zoomed-in positions of the heatmaps on the top. **(E)** We focused on mutations that are present in the DGRP data. We reported the distribution of the SSE scores predicted by the models **(F)** SSE distribution for DGRP positions. Each bar shows one location and the color indicates the TF binding site in the location of the SNP. Each dot is the average SSE score of one cluster from figure 1 D. **(G)** Each point in the scatter plot represents an individual within the DGRP data. The color of the points represents overlapping individuals. The x-axis is the SNP with the largest effect that the individual has, and the y-axis is the expected prediction of models when we use the entire enhancer of the individual. The black line represents the y=x. The individuals are from the DGRP data. The plot shows that most of the individuals enhancers have their largest effect SNP compensated by other SNPs within the enhancer.

We noted above examples of single nucleotide mutations that could potentially cause a large change in gene expression, although one might expect the more impactful mutations to be avoided in a population, given that our analysis focuses on an early developmental enhancer (42). We compared these impactful mutations to the allele frequencies of SNPs recorded in a population of 205 lines, as per the Drosophila Genome Reference Panel (DGRP), and noted that these mutations are indeed absent from this population. To further explore the relationship between predicted functional impact and allele frequencies, we examined the 70 positions in the 1416 bp-long *ind* enhancer that are polymorphic in the DGRP population (Figure 2E). Eleven of the 70 SNPs fall within the annotated binding sites (marked by ‘*’ in Figure 2E), and another 7 SNPs give rise to new weak binding sites. In total, 18 of the 70 SNPs may cause changes to binding site strengths and thus to *ind* expression (Supplementary Figure 1D). We used the filtered ensemble to predict the effect of each of the 18 SNP positions separately (Figure 2F and Supplementary Figure 1D), and noted that the maximum of these predicted effects – SSE of 0.15 for position 309 – is relatively small compared to the largest predicted effect (SSE of 0.35 for position 1198, Figure 2D and Supplementary Figures 1A,C) among all possible mutations, suggesting that high impact mutations are avoided in the population, as expected.

Figure 2F and Supplementary Figure 1D also reveal that for several of these 18 SNP positions different models in the ensemble make mutually inconsistent predictions (high versus low effect). Such variance in predicted effects points to ambiguities in underlying mechanisms, and offers candidates for experimental testing: data on the true impact of a SNP with ambiguous effect should help constrain the filtered ensemble further and narrow down the viable groups of models further.

### Evidence of compensatory mutations in individual enhancers

Each line in the DGRP population may manifest zero, one or more of the above 18 SNPs, and the net effect of multiple alternative alleles present in a line may not be the sum of their individual effects, i.e., there may be compensatory effects from multiple mutations in an individual enhancer. To investigate this, we next used the filtered ensemble to predict the expression profile of each line’s enhancer-level genotype and compared the effect (SSE between this prediction and wild-type expression) to the largest effect from a single mutation carried by the line (Figure 2G). We noted that a great majority (82 %) of lines exhibited signs of compensatory mutations (points below the y=x line in Figure 2G), and that almost all lines were predicted to have an *ind* expression profile very similar to the wild-type profile (Supplementary Figure 2A). We also generated a large number of synthetic genotypes (see Methods) that harbor zero, one or more of the 18 SNPs while preserving allele frequencies of each SNP, and repeated the above exercise (Supplementary Figure 2B, C) to obtain a null distribution of the compensatory effect. The fraction of lines that exhibit compensatory mutations in the real population (82%, as noted above) was found to be significantly higher than that in the empirically estimated null distribution (52%) (p-value < 9.62 *10E-22), substantiating the observation of compensatory mutations within lines.

### Experimental testing of selected polymorphisms identifies a single mechanism

We identified (above) several single-nucleotide mutations, present in the population or otherwise, for which the filtered ensemble predicts a large average effect on *ind* expression or exhibits a high degree of uncertainty. We used *in vivo* reporter assays to test the expression pattern driven by three such variants (called “construct1”, “construct2”, “construct3” repecively and explained belo) of the *ind* enhancer.

The first experiment (“construct1”) was designed to test the single mutation that has the greatest predicted effect, averaged over the ensemble. This mutation (Figure 3A), a T -> G change at position 1198 in the enhancer, impacts a crucial residue in a high affinity VND binding site, which might result in ventral de-repression of the enhancer, i.e., in its ventral border expanding. The mutation is not seen in the DGRP population, but was selected for the high average and moderate variance in predicted effect. In particular, while the mean prediction of the ensemble of models was a significant ventral de-repression, a subgroup of models in the ensemble also predicted no change in expression, indicating ambiguity in the ensemble.

**Figure 3.**
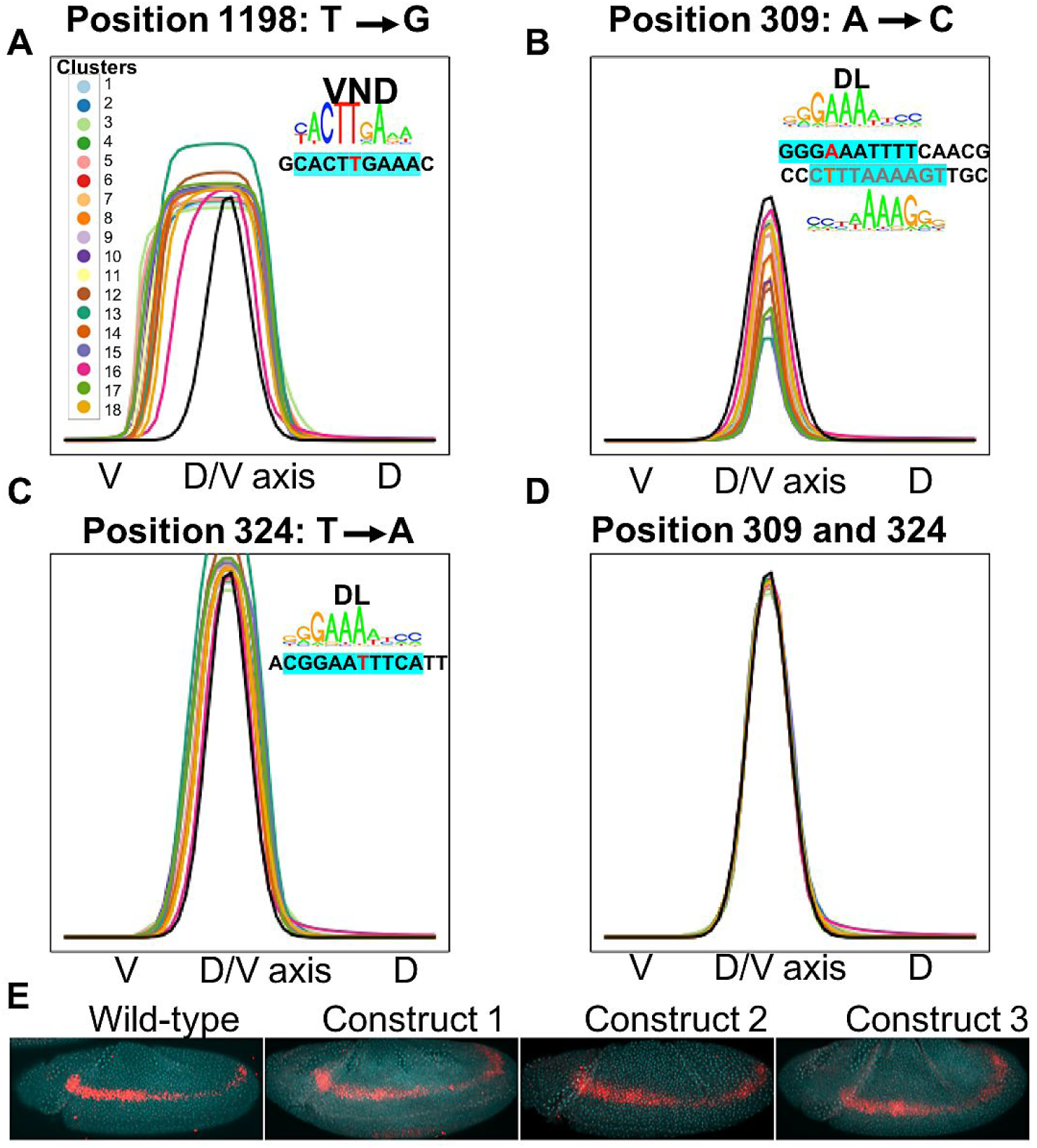
Polymorphism candidates for in vivo testing. **(A)** Different models predict the impact of SNPs on gene expression profile of ind enhancer. Each cluster or group of models are marked in a color and the plotted expression profile is the average expression predicted from models in that cluster. This SNP does not exist in the wild-type population and it hits a perfect VND site. **(B)** The SNP selected in this part exists in the wild-type DGRP data. There are two weak DL binding sites overlapping this position and they are conserved across drosophila species. Multiple groups of models predict a large impact. **(C)** Individuals with the SNP in part B tend to have this SNP that we suspect to offset the effect of losing DL binding site **(D)** Compensatory effect of two SNPs in part B and C. **(E)** Embryos from flies expressing lacZ under the control of either WT or variant enhancers stained with probes for lacZ mRNA (red) and DAPI (cyan).

The second experiment (“construct2”) was designed to test a mutation with a high uncertainty, i.e., large variance among predicted effects from different groups of models. This variant (Figure 3B), an A -> C change at position 309 in the enhancer, is predicted to reduce the binding strengths of two overlapping binding sites of the activator DL. The predicted impact of this mutation, which is seen in 6.1% of the DGRP lines, is a ∼25% reduction in the peak height (maximum expression level) on average, but there are groups of models that predict over 50% reduction and those that predict almost no change in expression. Interestingly, the same group of models (cluster 16, Figures 3A,B) predicted the smallest change for both of these enhancer variants.

The third construct (“construct3”) tested harbors two single nucleotide differences from the wild-type, and was predicted unanimously by all models in the ensemble to have no effect (Figure 3D). The two variants, which are present together in six DGRP lines, include the A->C change at position 309, introduced in the previous paragraph, that should decrease DL binding strength, and another mutation, a T->A change at position 324, also located within a DL binding site and predicted to increase DL binding strength. Together these two changes are predicted by several groups of models to compensate each other, while at least one group predicts neither to impact expression (see Supplementary Figure 3).

We tested the three above variants of the *ind* enhancer, as well as the wild-type enhancer, through reporter transgenic embryos and confocal laser scanning microscopy (Figure 3E, see Methods for details). All three variant enhancers were found to recapitulate the wild-type expression pattern of *ind.* There is exactly one group of models among 18 distinct groups in the filtered ensemble whose predictions are consistent with these new experimental data (Supplementary figure 4). In other words, tests of three carefully chosen variant enhancers allowed us to dramatically reduce the space of mechanistic explanations (see Figure 1E) to that represented by a tightly clustered group of models, ostensibly representing a single mechanistic explanation of *ind* regulation. We refer to this group of models as the “final ensemble”.

### Final ensemble is consistent with orthologous enhancers

Orthologs of the *D. melanogaster ind* enhancer from other Drosophila species are expected to drive similar expression patterns, given the key role played by this gene in early embryonic development. Under this assumption (also made elsewhere, e.g., (43–45)), orthologs provide an opportunity to cross-validate models of enhancer function: accurate models when applied to an ortholog may be reasonably expected to predict an expression pattern similar to the known *D. melanogaster* pattern. We therefore predicted the expression pattern driven by 10 different orthologs of the *D. melanogaster* enhancer, using the final ensemble alone or using every group of models in the filtered ensemble (Figure 4). We noted that the final ensemble makes accurate predictions for the majority of orthologs (Figure 4B, D, F, H, L, N), and provides more accurate predictions on the entire collection of orthologs, compared to other groups of models (Figure 4U. For instance, for the most diverged ortholog – that from *D. mojavensis* – the final ensemble is the only group of models that predicts expression in the correct location (Figure 4S, T).

**Figure 4.**
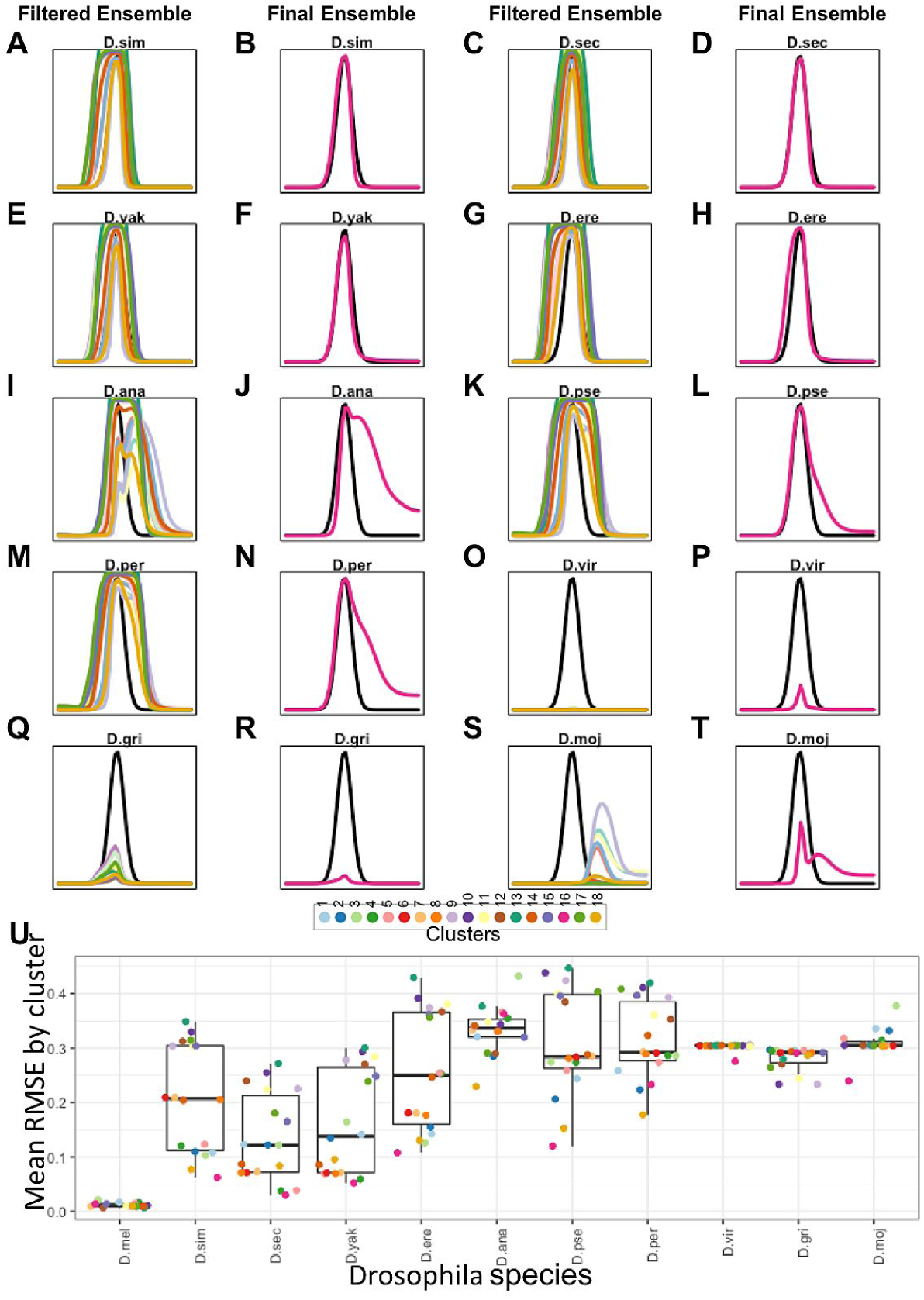
Cross validation with ortholog enhancers. For each ortholog enhancer sequence, we predicted the expression profile using models trained on the melanogaster. Each color shows the average expression predicted by models that cluster in the same group. The magenta curves in the second and fourth columns are the group of model that predicted the effect of polymorphism correctly (The final ensemble). We observe that for distant enhancers, the magenta curve predicts the expression in the correct position along the D/V axis while other groups of models misplace the peak of expression (moj) or cannot produce any expression (vir). **(A-B)** D.sim: Drosophila Simulans. **(C-D)**. D.sec: Drosophila Sechellia **(E-F)** D.yak: Yakuba **(G-H)** D.ere: Drosiphila Erecta. **(I-J)** D.ana: Drosiphila Ananassae. **(K-L)** D.pse: Drosiphila. **(M-N)** D.per: Drosiphila Persimilis. **(O-P)** D.vir: Drosiphila Virilis. **(Q-R)** D.gre: Drosiphila Grimshawi. **(S-T)** D.moj: Drosiphila Mojavensis. **(U)** Expected rmse scores between the expression predicted by the models and the wildtype ind expression for each species colored by the cluster.

### Final ensemble makes accurate predictions on variants of *rhomboid* enhancer

In another attempt to test if the final ensemble is more accurate compared to other groups of models in the filtered ensemble, we compared its predictions on the wild-type enhancer of a different neuroectodermal gene, *rhomboid (rho)*. The *rho* gene has an expression pattern similar to that of the *ind* gene and its enhancer is well studied; in fact, Sayal et al. (19) experimentally characterized the expression pattern driven by this enhancer as well as 37 synthetic variants thereof. Since the *ind* enhancer (subject of our modeling above) and the *rho* enhancer have similar expression patterns (outputs) and share regulators (inputs), we sought to cross-validate our models, trained with *ind* data, on the *rho* enhancer and its variants (19).

The *rho* enhancer is known to be controlled by two activators – DL and TWI – and one repressor, SNA. While DL and SNA were among the TFs included in the models of *ind* above, TWI was not, and as a result the trained models are not capable of predicting *rho* expression. To address this, we performed partial optimization of parameters on the *rho* data set (37 synthetic constructs) from Sayal et al (19). In particular, we considered each model trained on *ind* data (previous sections), utilized the trained values of four of its parameters that are shared between *ind* and *rho* models without further modification, but trained four additional parameters unique to *rho* on the *rho* data set (see Methods). As a result, each model in the filtered ensemble from above gives rise to a model for the *rho* data, with four unchanged parameters and four newly trained parameters (Figure 5A). The accuracy of the resulting model on each of the *rho* enhancers (wild type and its variants) was then assessed using SSE score. We noted that not all models in the filtered ensemble led to models capable of explaining the *rho* data set; rather, only models belonging to 10 of the 18 groups of models in the ensemble could, upon training of the additional parameters, provide fits better than a modest threshold of SSE = 0.1. The prediction of each of these 10 groups of models for the wild-type *rho* enhancer is shown in Figure 5B.

**Figure 5.**
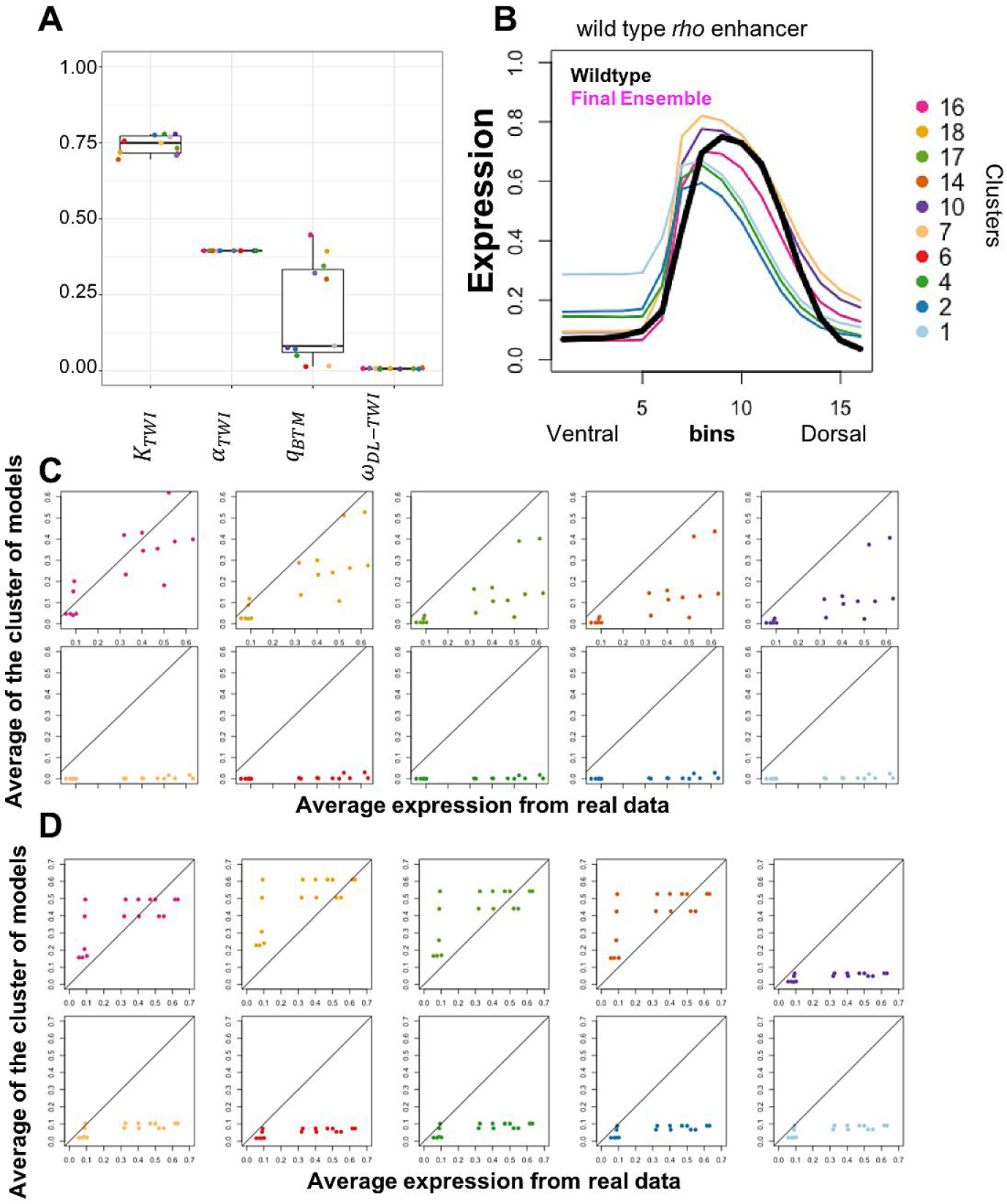
Cross validation with rho enhancers. We used eight of the parameters of TFs present in ind models and trained four extra parameters to model thirty-eight different rho enhancer constructs. Then we filtered models with good SSE scores. 10 out of 18 clusters from ind models survive the filtering (including the final cluster that survived the SNP filters). **(A)** four new parameters are trained on wildtype ‘rho’ enhancer starting from the parameters trained on ‘ind’. The distribution of the new parameters is shown in the box plots where each dot represents a group of models in the original ‘ind’ enhancer. **(B)** wildtype expression profile of ‘rho’ enhancer predicted by the models. The magenta is the final ensemble **(C)** Average expression in the bins near the peak of the rho enhancer along the D/V axis (bins 8-12). The x-axis is the average expression from the real data and the y-axis is the model predicted average expression. Each dot is a construct in the dataset. Each panel is the from a group of models colored by the clusters in B. The top left panel is the final ensemble. **(D)** Similar to C, the average expression is computed in the first 5 bins (bins 1 to 5). The y-axis is the model predicted effect and the x-axis is the real data.

We next examined predictions of the above models on 26 enhancers that differed from the wild-type *rho* enhancer in that the peak expression driven by these variant enhancers is significantly lower than that of the wild-type enhancer (Supplementary Figure 5). We assessed how well the models in each group from the filtered ensemble capture this phenomenon: for each group of models we computed the average predicted expression in the peak expression region and compared it to the true (experimentally measured) expression in that region (Figure 5C). It was visually clear that models from the final ensemble (top left panel in Figure 5C) captured the reduced peak expression levels of these 26 variant enhancers better than all other groups of models of the filtered ensemble. We similarly examined predictions on the 9 enhancers that differ from the *rho* enhancer in a clear de-repression in the ventral-most region of the embryo (Supplementary Figure 6). For these enhancers, we computed the predicted expression, from each group of models, in this spatial region and compared it to the experimentally observed expression in the region (Figure 5D). Visual inspection reveals that the models originating in the final ensemble capture the phenomenon of ventral derepression better than six of the other groups of models and at least as well as the remaining three groups in the filtered ensemble. In summary, the final ensemble identified above based on our experimental assessment of mutation effects proved to be far more accurate than competing groups of models, in terms of its ability to generalize to a new data set comprising a related but distinct enhancer (the *rho* enhancer) and its variants.

## DISCUSSION

A major open problem today is how DNA sequence variations, e.g., single-nucleotide polymorphisms (SNPs), lead to phenotypic differences among individuals. A popular approach is to find polymorphisms that are statistically correlated with the phenotype, as in genome-wide association studies (GWAS) (4), family-based association tests (46), and expression quantitative trait loci (eQTLs) (47, 48) for phenotype-related genes. However, statistically identified variations may not be functionally related to phenotypes (49), due to a variety of factors including linkage disequilibrium (LD) and redundancy of genetic systems. This problem is particularly pronounced in the case of non-coding variations, which form the majority of GWAS findings (50) and function by influencing gene regulation. Accurate contextual or mechanistic information about non-coding variations can help us pinpoint those that are causally related to the phenotype (5, 31). The work presented here is a step in this direction, and provides an example of how detailed mechanistic models of the sequence-to-expression relationship encoded by an enhancer may help us predict the effects of non-coding variations. This in turn can lead to better prioritization of phenotype-related variants and also provide mechanistic explanation of their effects.

In recent years, various machine learning-based methods such as gkm-SVM (25) and DeepSEA (26) have been proposed for modeling the sequence-function relationship encoded throughout the non-coding genome. These have been successful in predicting the impact of variants on epigenomic states such as DNA accessibility and TF-DNA binding (25, 26), although some reports indicate there is significant room for improvement in their accuracy (51) There is also evidence that these machine learning methods can help identify eQTLs and disease-related variations. The thermodynamics-based modeling approach used in our work offers a complementary approach to variant interpretation, and while it is far less scalable than the ML-based methods it is more mechanistically grounded and potentially more precise. It is also possible that similar models, once trained on high throughput data such as those from massively parallel reporter assays (52), will provide mechanistic predictions about non-coding variations on a larger scale than the current work. In a simple illustration of how mechanistic models can be useful at scale, Xie et al. (53) showed that motif-based biophysics-inspired models of TF-DNA binding predict SNPs that likely impact binding strength and lead to inter-individual variation in chemosensitivity.

The use of mechanistic quantitative models to examine polymorphisms, though uncommon today, is not entirely new. Gursky et al. (28) analyzed polymorphisms in Drosophila strains (the same collection as in our study) using the same sequence-to-expression model. They reported several valuable insights, including additive effect of multiple polymorphisms in individual genotypes and evidence of selective pressure at the level of combinations of SNPs. Our analysis has conceptual and methodological similarities to Gursky et al., with a few key differences. First, our approach recognizes that uncertainties in the model (values of parameters) can lead to ambiguities in polymorphism analysis, and our predictions are accompanied by estimates of the resulting uncertainty. More importantly, while Gursky et al. focused primarily using the sequence-to-expression model to reveal insights about a collection of polymorphisms, we focus more on functional analysis and use experimental assays of variant effects to refine the models, making them more precise and more ready for future applications.

Our work also underscores the value of model-based design of biological experiments. Two of the three enhancer variants that we tested were chosen because models based on prior data were ambiguous in their predictions regarding those variants. After we performed those experiments, the results led to a significant narrowing of the feasible models and this smaller feasible group of models was then shown to be more consistent with held-out data sets (based on orthologs of the ind enhancer as well as several synthetic variants of the rho enhancer) than the original broader ensemble of models. We hope that such iterative applications of modeling and experimental testing, with models furnishing candidates for experimentation and experimental results refining the models, will be more frequently adopted in future investigations.

## AUTHOR CONTRIBUTIONS

FK performed research; analyzed data; designed research; wrote the manuscript.

NS: performed research and wrote the manuscript

ND: performed research and wrote the manuscript.

GJ: designed research and wrote the manuscript

SS: designed research and wrote the manuscript

## ACKNOWLEDGMENTS

The authors would like to thank professor Stanislav Shvartsman for his valuable input and feedback on this manuscript. SS and FK were supported by the National Institutes of Health (R01GM114341, R35GM131819). NS and GJ were supported by research grants from the Spanish Government (BFU2017-87244-P) and Generalitat de Catalunya (2017 SGR 475).

## REFERENCES

1. Hamburg, M.A., and F.S. Collins. 2010. The Path to Personalized Medicine - Perspective. N. Engl. J. Med. 363: 301–304.

2. Haraksingh, R.R., and M.P. Snyder. 2013. Impacts of variation in the human genome on gene regulation. J. Mol. Biol. 425: 3970–3977.

3. Morloy, M., C.M. Molony, T.M. Weber, J.L. Devlin, K.G. Ewens, R.S. Spielman, and V.G. Cheung. 2004. Genetic analysis of genome-wide variation in human gene expression. Nature. 430: 743.

4. Zhang, F., and J.R. Lupski. 2015. Non-coding genetic variants in human disease. Hum. Mol. Genet. 24: R102–R110.

5. Maurano, M.T., R. Humbert, E. Rynes, R.E. Thurman, E. Haugen, H. Wang, A.P. Reynolds, R. Sandstrom, H. Qu, J. Brody, A. Shafer, F. Neri, K. Lee, T. Kutyavin, S. Stehling-Sun, A.K. Johnson, T.K. Canfield, E. Giste, M. Diegel, D. Bates, R.S. Hansen, S. Neph, P.J. Sabo, S. Heimfeld, A. Raubitschek, S. Ziegler, C. Cotsapas, N. Sotoodehnia, I. Glass, S.R. Sunyaev, R. Kaul, and J.A. Stamatoyannopoulos. 2012. Systematic localization of common disease-associated variation in regulatory DNA. Science (80-.). 337: 1190–1195.

6. Ward, L.D., and M. Kellis. 2012. Interpreting noncoding genetic variation in complex traits and human disease. Nat. Biotechnol. 30: 1095–106.

7. Maston, G.A., S.K. Evans, and M.R. Green. 2006. Transcriptional Regulatory Elements in the Human Genome. Annu. Rev. Genomics Hum. Genet. 7: 29–59.

8. Dunham, I., A. Kundaje, S.F. Aldred, P.J. Collins, C.A. Davis, F. Doyle, C.B. Epstein, S. Frietze, J. Harrow, R. Kaul, J. Khatun, B.R. Lajoie, S.G. Landt, B.K. Lee, F. Pauli, K.R. Rosenbloom, P. Sabo, A. Safi, A. Sanyal, N. Shoresh, J.M. Simon, L. Song, N.D. Trinklein, R.C. Altshuler, E. Birney, J.B. Brown, C. Cheng, S. Djebali, X. Dong, J. Ernst, T.S. Furey, M. Gerstein, B. Giardine, M. Greven, R.C. Hardison, R.S. Harris, J. Herrero, M.M. Hoffman, S. Iyer, M. Kellis, P. Kheradpour, T. Lassmann, Q. Li, X. Lin, G.K. Marinov, A. Merkel, A. Mortazavi, S.C.J. Parker, T.E. Reddy, J. Rozowsky, F. Schlesinger, R.E. Thurman, J. Wang, L.D. Ward, T.W. Whitfield, S.P. Wilder, W. Wu, H.S. Xi, K.Y. Yip, J. Zhuang, B.E. Bernstein, E.D. Green, C. Gunter, M. Snyder, M.J. Pazin, R.F. Lowdon, L.A.L. Dillon, L.B. Adams, C.J. Kelly, J. Zhang, J.R. Wexler, P.J. Good, E.A. Feingold, G.E. Crawford, J. Dekker, L. Elnitski, P.J. Farnham, M.C. Giddings, T.R. Gingeras, R. Guigó, T.J. Hubbard, W.J. Kent, J.D. Lieb, E.H. Margulies, R.M. Myers, J.A. Stamatoyannopoulos, S.A. Tenenbaum, Z. Weng, K.P. White, B. Wold, Y. Yu, J. Wrobel, B.A. Risk, H.P. Gunawardena, H.C. Kuiper, C.W. Maier, L. Xie, X. Chen, et al. 2012. ENCODE Project Consortium. An integrated encyclopedia of DNA elements in the human genome. Nature. 489: 57–74.

9. Korte, A., and A. Farlow. 2013. The advantages and limitations of trait analysis with GWAS: A review. Plant Methods..

10. Tak, Y.G., and P.J. Farnham. 2015. Making sense of GWAS: Using epigenomics and genome engineering to understand the functional relevance of SNPs in non-coding regions of the human genome. Epigenetics and Chromatin. 8: 57.

11. Pope, B.D., T. Ryba, V. Dileep, F. Yue, W. Wu, O. Denas, D.L. Vera, Y. Wang, R.S. Hansen, T.K. Canfield, R.E. Thurman, Y. Cheng, G. Gülsoy, J.H. Dennis, M.P. Snyder, J.A. Stamatoyannopoulos, J. Taylor, R.C. Hardison, T. Kahveci, B. Ren, and D.M. Gilbert. 2014. Topologically associating domains are stable units of replication-timing regulation. Nature. 515: 402.

12. Li, W., D. Notani, and M.G. Rosenfeld. 2016. Enhancers as non-coding RNA transcription units: Recent insights and future perspectives. Nat. Rev. Genet. 17: 207–223.

13. Visel, A., E.M. Rubin, and L.A. Pennacchio. 2009. Genomic views of distant-acting enhancers. Nature. 461: 199.

14. Kasowski, M., F. Grubert, C. Heffelfinger, M. Hariharan, A. Asabere, S.M. Waszak, L. Habegger, J. Rozowsky, M. Shi, A.E. Urban, M.Y. Hong, K.J. Karczewski, W. Huber, S.M. Weissman, M.B. Gerstein, J.O. Korbel, and M. Snyder. 2010. Variation in transcription factor binding among humans. Science (80-.). 328: 232–235.

15. Cusanovich, D.A., B. Pavlovic, J.K. Pritchard, and Y. Gilad. 2014. The Functional Consequences of Variation in Transcription Factor Binding. PLoS Genet. 10: e1004226.

16. Segal, E., T. Raveh-Sadka, M. Schroeder, U. Unnerstall, and U. Gaul. 2008. Predicting expression patterns from regulatory sequence in Drosophila segmentation. Nature. 451: 535–540.

17. Janssens, H., S. Hou, J. Jaeger, A.R. Kim, E. Myasnikova, D. Sharp, and J. Reinitz. 2006. Quantitative and predictive model of transcriptional control of the Drosophila melanogaster even skipped gene. Nat. Genet. 38: 1159–1165.

18. Zinzen, R.P., K. Senger, M. Levine, and D. Papatsenko. 2006. Computational Models for Neurogenic Gene Expression in the Drosophila Embryo. Curr. Biol. 16: 1358–1365.

19. Sayal, R., J.M. Dresch, I. Pushel, B.R. Taylor, and D.N. Arnosti. 2016. Quantitative perturbation-based analysis of gene expression predicts enhancer activity in early Drosophila embryo. Elife. 5: 1–25.

20. White, M.A., D.S. Parker, S. Barolo, and B.A. Cohen. 2012. A model of spatially restricted transcription in opposing gradients of activators and repressors. Mol. Syst. Biol. 8.

21. Ahsendorf, T., F. Wong, R. Eils, and J. Gunawardena. 2014. A framework for modelling gene regulation which accommodates non-equilibrium mechanisms. BMC Biol. 12: 102.

22. He, X., M.A.H. Samee, C. Blatti, and S. Sinha. 2010. Thermodynamics-based models of transcriptional regulation by enhancers: The roles of synergistic activation, cooperative binding and short-range repression. PLoS Comput. Biol. 6: e1000935.

23. Kaplan, T., X.-Y. Li, P.J. Sabo, S. Thomas, J.A. Stamatoyannopoulos, M.D. Biggin, and M.B. Eisen. 2011. Quantitative Models of the Mechanisms That Control Genome-Wide Patterns of Transcription Factor Binding during Early Drosophila Development. PLoS Genet. 7: e1001290.

24. Ay, A., and D.N. Arnosti. 2011. Mathematical modeling of gene expression: a guide for the perplexed biologist. Crit. Rev. Biochem. Mol. Biol. 46: 137–151.

25. Lee, D., D.U. Gorkin, M. Baker, B.J. Strober, A.L. Asoni, A.S. Mccallion, and M.A. Beer. 2015. A method to predict the impact of regulatory variants from DNA sequence. Nat. Genet. 47: 955–961.

26. Zhou, J., and O.G. Troyanskaya. 2015. Predicting effects of noncoding variants with deep learning–based sequence model. Nat. Methods. 12: 931–934.

27. Samee, M.A.H., B. Lim, N. Samper, H. Lu, C.A. Rushlow, G. Jiménez, S.Y. Shvartsman, and S. Sinha. 2015. A Systematic Ensemble Approach to Thermodynamic Modeling of Gene Expression from Sequence Data. Cell Syst. 1: 396–407.

28. Gursky, V. V, K.N. Kozlov, I. V Kulakovskiy, A. Zubair, P. Marjoram, D.S. Lawrie, S. V Nuzhdin, and M.G. Samsonova. 2017. Translating natural genetic variation to gene expression in a computational model of the Drosophila gap gene regulatory network..

29. Khajouei, F., and S. Sinha. 2018. An information theoretic treatment of sequence-to-expression modeling. PLOS Comput. Biol. 14: e1006459.

30. Yáñez-Cuna, J.O., E.Z. Kvon, and A. Stark. 2013. Deciphering the transcriptional cis-regulatory code. Trends Genet. 29: 11–22.

31. Deplancke, B., D. Alpern, and V. Gardeux. 2016. The Genetics of Transcription Factor DNA Binding Variation. Cell. 166: 538–554.

32. Nien, C.Y., H.L. Liang, S. Butcher, Y. Sun, S. Fu, T. Gocha, N. Kirov, J.R. Manak, and C. Rushlow. 2011. Temporal coordination of gene networks by Zelda in the early Drosophila embryo. PLoS Genet. 7.

33. Garcia, M., and A. Stathopoulos. 2011. Lateral gene expression in Drosophila early embryos is supported by grainyhead-mediated activation and tiers of dorsally-localized repression. PLoS One. 6: e29172.

34. Ohlen, T. Von, and C.Q. Doe. 2000. Convergence of Dorsal, Dpp, and Egfr Signaling Pathways Subdivides the Drosophila Neuroectoderm into Three Dorsal-Ventral Columns. 372: 362–372.

35. McDonald, J.A., S. Holbrook, T. Isshiki, J. Weiss, C.Q. Doe, and D.M. Mellerick. 1998. Dorsoventral patterning in the Drosophila central nervous system: The vnd homeobox gene specifies ventral column identity. Genes Dev. 12: 3603–3612.

36. Lim, B., N. Samper, H. Lu, C. Rushlow, G. Jiménez, and S.Y. Shvartsman. 2013. Kinetics of gene derepression by ERK signaling. Proc. Natl. Acad. Sci. U. S. A. 110: 10330–5.

37. Bischof, J., M. Björklund, E. Furger, C. Schertel, J. Taipale, and K. Basler. 2012. A versatile platform for creating a comprehensive UAS-ORFeome library in Drosophila. Dev. 140: 2434–2442.

38. Bischof, J., R.K. Maeda, M. Hediger, F. Karch, and K. Basler. 2007. An optimized transgenesis system for Drosophila using germ-line-specific φC31 integrases. Proc. Natl. Acad. Sci. U. S. A. 104: 3312–3317.

39. Gertz, J., E.D. Siggia, and B.A. Cohen. 2009. Analysis of combinatorial cis-regulation in synthetic and genomic promoters. Nature. 457: 215–218.

40. Suleimenov, Y., A. Ay, M.A.H. Samee, J.M. Dresch, S. Sinha, and D.N. Arnosti. 2013. Global parameter estimation for thermodynamic models of transcriptional regulation. Methods. 62: 99–108.

41. Huang, W., A. Massouras, Y. Inoue, J. Peiffer, M. Ràmia, A.M. Tarone, L. Turlapati, T. Zichner, D. Zhu, R.F. Lyman, M.M. Magwire, K. Blankenburg, M.A. Carbone, K. Chang, L.L. Ellis, S. Fernandez, Y. Han, G. Highnam, C.E. Hjelmen, J.R. Jack, M. Javaid, J. Jayaseelan, D. Kalra, S. Lee, L. Lewis, M. Munidasa, F. Ongeri, S. Patel, L. Perales, A. Perez, L.L. Pu, S.M. Rollmann, R. Ruth, N. Saada, C. Warner, A. Williams, Y.Q. Wu, A. Yamamoto, Y. Zhang, Y. Zhu, R.R.H. Anholt, J.O. Korbel, D. Mittelman, D.M. Muzny, R.A. Gibbs, A. Barbadilla, J.S. Johnston, E.A. Stone, S. Richards, B. Deplancke, and T.F.C. Mackay. 2014. Natural variation in genome architecture among 205 Drosophila melanogaster Genetic Reference Panel lines. Genome Res. 24: 1193–1208.

42. Phinchongsakuldit, J., S. MacArthur, and J.F.Y. Brookfield. 2004. Evolution of Developmental Genes: Molecular Microevolution of Enhancer Sequences at the Ubx Locus in Drosophila, and Its Impact on Developmental Phenotypes. Mol. Biol. Evol. 21: 348–363.

43. Zeitlinger, J., R.P. Zinzen, A. Stark, M. Kellis, H. Zhang, R.A. Young, and M. Levine. 2007. Whole-genome ChIP-chip analysis of Dorsal, Twist, and Snail suggests integration of diverse patterning processes in the Drosophila embryo. Genes Dev. 21: 385–390.

44. Werner, T., A. Hammer, M. Wahlbuhl, M.R. Bösl, and M. Wegner. 2007. Multiple conserved regulatory elements with overlapping functions determine Sox10 expression in mouse embryogenesis. Nucleic Acids Res. 35: 6526–6538.

45. Hong, J.W., D.A. Hendrix, and M.S. Levine. 2008. Shadow enhancers as a source of evolutionary novelty. Science (80-.). 321: 1314–1314.

46. Horn, S., A. Figl, P.S. Rachakonda, C. Fischer, A. Sucker, A. Gast, S. Kadel, I. Moll, E. Nagore, K. Hemminki, D. Schadendorf, and R. Kumar. 2013. TERT promoter mutations in familial and sporadic melanoma. Science (80-.). 339: 959–961.

47. Cookson, W., L. Liang, G. Abecasis, M. Moffatt, and M. Lathrop. 2009. Mapping complex disease traits with global gene expression. Nat. Rev. Genet. 10: 184.

48. Croteau-Chonka, D.C., A.J. Rogers, T. Raj, M.J. McGeachie, W. Qiu, J.P. Ziniti, B.J. Stubbs, L. Liang, F.D. Martinez, R.C. Strunk, R.F. Lemanske, A.H. Liu, B.E. Stranger, V.J. Carey, and B.A. Raby. 2015. Expression quantitative trait loci information improves predictive modeling of disease relevance of non-coding genetic variation. PLoS One. 10: e0140758.

49. Boyle, E.A., Y.I. Li, and J.K. Pritchard. 2017. An Expanded View of Complex Traits: From Polygenic to Omnigenic. Cell. 169: 1177–1186.

50. Paul, D., N. Soranzo, and S. Beck. 2014. Functional interpretation of non-coding sequence variation: Concepts and challenges. BioEssays. 6: 191–199.

51. Wagih, O., D. Merico, A. Delong, and B.J. Frey. 2018. Allele-specific transcription factor binding as a benchmark for assessing variant impact predictors. bioRxiv.: 253427.

52. Mogno, I., J.C. Kwasnieski, and B.A. Cohen. 2013. Massively parallel synthetic promoter assays reveal the in vivo effects of binding site variants. Genome Res. 23: 1908–1915.

53. Xie, X., C. Hanson, and S. Sinha. 2019. Mechanistic interpretation of non-coding variants for discovering transcriptional regulators of drug response. BMC Biol. 17.

